# Rational use of Episodic and Working Memory: A Normative Account of Prospective Memory

**DOI:** 10.1101/580324

**Authors:** Ida Momennejad, Jarrod Lewis-Peacock, Kennneth A. Norman, Jonathan Cohen, Satinder Singh, Richard L. Lewis

**Affiliations:** Department of Biomedical Engineering, Columbia University; Department of Psychology, University of Texas, Austin, TX; Department of Psychology, Princeton Neuroscience Institute, Princeton University; Computer Science & Engineering Division, Department of Elec. Eng. & Computer Science, University of Michigan; Department of Psychology, Weinberg Institute for Cognitive Science, University of Michigan

## Abstract

Humans often simultaneously pursue multiple plans at different time scales. The successful realization of non-immediate plans (e.g., post package after work) requires keeping track of a future plan while accomplishing other intermediate tasks (e.g., write a paper), a capacity known as *prospective memory*. This capacity requires the integration of noisy evidence from perceptual input with evidence from short-term working memory (WM) and longer-term or episodic memory (LTM/EM). Here we formulate a set of dual-task problems in empirical studies of prospective memory as problems of computational rationality, and ask how a rational model should exploit noisy perception and memory to maximize payoffs. The model combines reinforcement learning (optimal action selection) with evidence accumulation (optimal inference) in order to derive good decision parameters for optimal task performance (i.e., performing an ongoing task while monitoring for a cue that triggers executing a second prospective task). We compare model behavior to key accuracy and reaction time phenomena in human performance. Thus, we offer a normative approach to theorizing and modeling these phenomena without assumptions about mechanisms of attention or retrieval. This approach can be extended to study meta-parameters governing the boundedly rational use of memory in planned action in health, as well as compensatory mnemonic strategies that may be rational responses to disturbances of these mechanisms in neuropsychiatric disorders.

## 1 Introduction

The execution of cognitive control in the service of goal-directed behavior is intimately bound to the use of memory. The immediate pursuit of a goal is presumed to rely on control representations actively maintained in working memory (Anderson, 1983; Miller and Cohen, 2001). In contrast, pursuit of a future goal requires that it be encoded in longer-term storage, and retrieved at the appropriate time (Cohen et al., 1996; Gollwitzer and Brandstätter, 1997). This capacity is often referred to as prospective memory (PM). Prospective memory experiments are typically designed to study the interaction between performing an ongoing task (e.g., a day’s work) and the flexible encoding, retrieval, and realization of a prospective task (e.g., post a package). In event-based prospective memory, the prospective task must be executed as soon as the relevant target is perceived (e.g., the post office) (Einstein and McDaniel, 2005; Einstein et al., 2005). In time-based prospective memory (e.g., take out the cookies from the oven in 30 minutes), internal or external time keeping determines ’when’ to execute the future plan (Momennejad and Haynes, 2012).

PM success relies on rational use of memory and attentional processes. For instance, if you plan to post a package after a workday, you cannot rely on actively maintaining and rehearsing this plan in working memory all day, as this would interfere with your performance at work. At the same time, you must ensure that it is remembered in the face of the day’s workload, and reliably retrieved at the end of the day. The multiprocess view of PM (Einstein and McDaniel, 2005; Einstein et al., 2005) suggests that the successful realization of prospective memory tasks relies on a dynamic interaction between two categories of processes: effortful and controlled attentional processes to monitor for targets in the environment (e.g., active monitoring of the environment and mental rehearsal of the planned action), and spontaneous retrieval processes (i.e., spontaneously remembering to execute the plan once the target, say post office, is detected).

Here, we adopt the multiprocess framework and propose that the functioning of prospective memory relies on the *adaptive* use of working memory (WM), long-term/episodic memory (EM), and perception. In this model, top-down or monitoring processes of the multiprocess framework are operationalized as strategies that rely on working memory, while bottom-up or spontaneous retrieval processes are strategies that rely on long-term or episodic memory. In our account the weighing of the two sources of memory in the service of action selection is not hand-tuned, but derived by a normative model seeking to simultaneously maximize reward in both the ongoing and prospective tasks. This is accomplished by selecting actions according to an optimal policy (reinforcement learning) conditioned on a Bayesian integration of perceptoin and memory (inference).

It is important to highlight that our model is not a mechanistic process model of PM; rather, our contribution is to show that key findings in the prospective memory literature, and more broadly multi-tasking, can be explained by considering how to normatively reconcile multiple sources of memory and perceptual evidence. Indeed, the novel and perhaps surprising implication of our model is that a fairly detailed account of human behavioral performance (accuracy and reaction times) is possible using quite abstract assumptions about memory and perception, and an assumption of rationality (Lewis et al., 2014).

The model makes minimal, abstract assumptions about the properties of three noisy component subsystems: WM, LTM/EM, and perception. We explore the implications of these assumptions for prospective memory by asking how a rational model would use these components to adaptively respond to the demands of a variety of specific experimental task settings. At the heart of the model is the optimal integration of noisy information from the three components over time, and the optimal selection of external task response actions and internal memory actions to maximize rewards.

Many of the interesting empirical signatures of prospective memory concern effects on both accuracy and speed of dualtask performance, so the model must incorporate processing dynamics. We model processing dynamics (in discrete time) as the sequential accumulation and optimal integration of evidence from perception and memory, and the optimal selection of actions that either wait and sample more evidence, respond to the ongoing task, or respond to a possible prospective memory target. As we will see below, the formal statement of what the model should do is mathematically simple. But the multiple sources of evidence (two memory stores plus perceptual sampling) and multiple possible actions in the dual-task setting (four) give rise to the following significant technical challenges for computing the optimal policy. First, even the relatively simple assumptions about memory and perception noise that we adopt here preclude an analytic solution to the optimal Bayesian integration of perception and memory. Second, the dual task structure requiring a selection among three task actions (plus a wait-and-continue-sampling action) precludes the use of classic sampling models such as drift-diffusion or the Sequential Probability Ratio Test, which are formulated for the selection among two choices conditioned on one decision variable.

We address these technical challenges by using a Monte Carlo technique to provide good approximations of the Bayesian integration, and by using reinforcement learning (RL) algorithms to compute a good approximation of the optimal action selection policy. As such, we do not hand-tune control parameters, wait times, or thresholds to fit empirical data; rather, they emerge from the RL policy optimization, and without any explicit thresholds on decision variables. The optimality assumptions provide a powerful analytic basis for reducing the space of considered strategies and methods of noisy information integration. Although the mechanistic assumptions about component subsystems are very simple, the model generates a rich set of predictions for error rates and reaction times that provide good accounts of key empirical phenomena in the prospective memory literature.

The paper has the following structure. In §2 we review key aspects of the experimental literature, and identify experimental paradigms that express a canonical set of phenomena, summarized in §2.2. In §3 we formally specify the normative model. We then report in §4 the results of simulations that test the model’s account of the key behavioral phenomena. We discuss the nature of the explanations that the model provides, novel predictions by the model that remain to be empirically tested, as well as aspects of the empirical data which are not well-fit by the model. We conclude with a summary of the model’s strengths and caveats, and a discussion of the prospects for applying the modeling method to other domains that involve the adaptive integration of perception with multiple memory systems.

## 2 An Empirical Paradigm for Prospective Memory and Key Findings

Many prospective memory experiments are variations of an *event-based* experimental paradigm originally proposed by Einstein and colleagues (Brewer et al., 2011; Einstein et al., 2005; Scullin et al., 2012). In this section we first describe the experimental design, then summarize key phenomena of interest emerging in human performance on these tasks.

### 2.1 Event-based Prospective memory: A Dual-Tasking Paradigm

In event-based prospective memory, the occurrence of an event (e.g., spotting the post office) triggers a switch from an ongoing task (e.g., talking on the phone) to a planned action (e.g., posting a package). There are two components to event-based experimental paradigms: an *ongoing task (OG)* that demands the majority of responses, and a *prospective memory task (PM)* that demands a response only when a relatively infrequent target probe or event occurs, referred to as the *PM target*.

Figure 1 illustrates an example of this canonical experimental paradigm for testing prospective memory. As the original paradigm, the ongoing task is to judge whether the word on the right matches the category presented on the left (e.g., on trial 1, cat is an ANIMAL, a match, hence the correct response is yes). Participants performed this task on its own to set the baseline performance on the ongoing task alone (baseline OG, or No-PM trials). In the prospective memory condition, participants were required to give a prospective memory response by pressing another button whenever the PM target (e.g., the syllable ’tor’ in this case) appeared on screen (e.g., on the 4th trial, tortoise’ is a PM target). The instructions indicated the level of priority or emphasis of the PM task relative to the ongoing task. For instance, high PM emphasis was instruction as follows: It is very important that you consider your main goal in this section to find absolutely every occurrence of the target item. In the high PM emphasis condition participants were to prioritize their attentional and memory resources to increase PM performance, even at the cost of ongoing task performance. Moderate emphasis instructions indicated a more relaxed prioritization.

**Figure 1:**
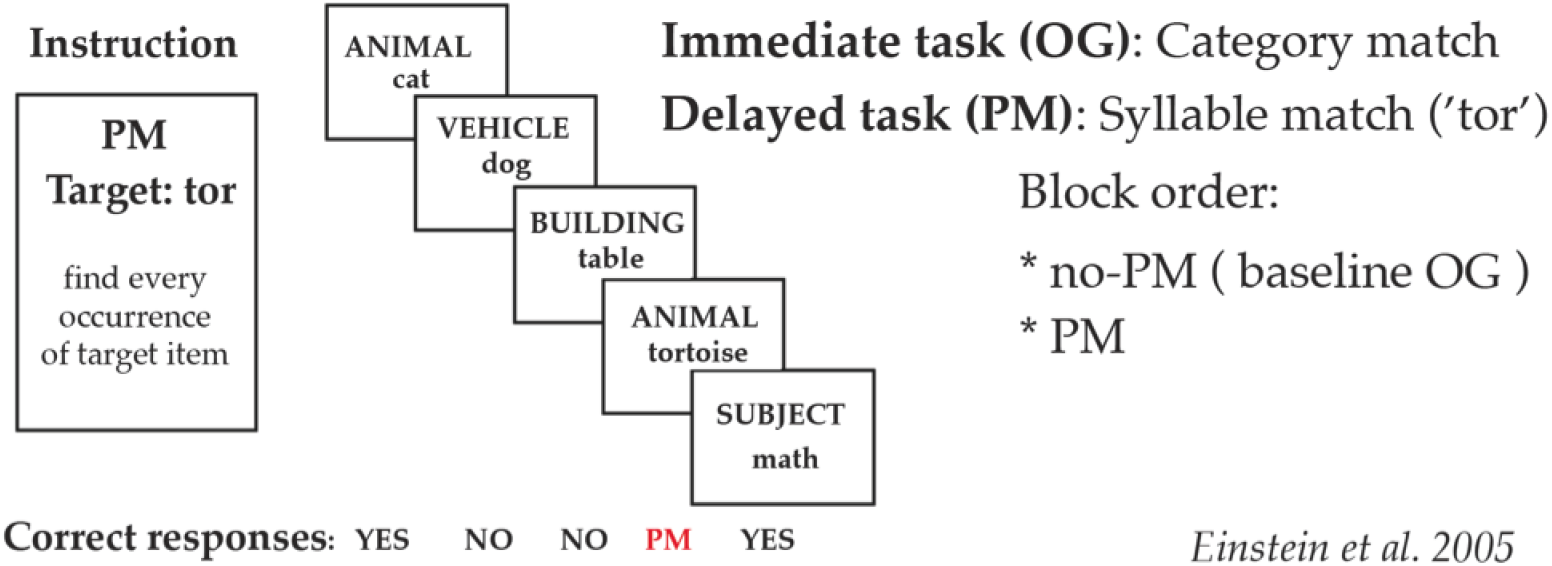
An example of an *event-based* prospective memory (PM) experimental paradigm (Einstein et al., 2005). The paradigm consists of a sequence of trials which have an ongoing task and a prospective memory target (PM target). A given trial begins with an instruction, indicating the prospective memory target, which was a different word or syllable on different trials. Each trial consists of a sequence of ongoing task (OG) stimuli requiring a category match judgment between two words on the screen. The PM target in the example instruction is the syllable *tor*, which here appears in the word *tortoise* in the fourth stimulus of the trial. The fourth stimulus thus requires a PM response. Here we have indicated correct responses to each stimulus below it for clarification. On half the trials the instruction emphasized the high importance of prospective memory accuracy (e.g., *It is very important that you consider your main goal in this section to find absolutely every occurrence of the target item*.). On the other half, the PM instruction was of moderate importance: *we have a secondary interest in your ability to remember to perform an action in the future*. On *no-PM* trials participants are instructed to perform only the ongoing task, setting a behavioral baseline for comparing reaction times to the ongoing task across conditions.

Next we review key findings from experiments that use variations of this experimental paradigm.

### 2.2 Key Phenomena in Event-based Prospective Memory

Prospective memory effects are generally measured and reported in terms of the influence of experimental conditions on two behavioral measures: accuracy of task performance and response times. Many PM studies have used variations of the experimental PM paradigm described in above, so we refer to this paradigm in the reported phenomena below. Studies commonly report an *OG cost* measure, which is the extent to which responses to the ongoing task are slowed as a function of PM-related experimental manipulations. Prospective memory performance itself is often reported in terms of *PM hit rate*, which measures the proportion of times a participant successfully detects and responds to a PM target. Below, we review four key classes of PM phenomena consistently reported in the literature. These key phenomena serve as the target for the behavior of the model described in the next section.

#### Effects of target focality

A PM target is *focal* when the OG and PM tasks both require attention to similar features of stimuli. For instance, a PM target is focal if the ongoing task is a word match judgment and the prospective task is a particular word (e.g., ’tortoise’). An example of non-focal PM is when the OG task is to judge word match but the PM task is to respond to the font color or a particular syllable (e.g., red font or any word with the syllable ’tor’).

The main effect of focality is that participants are more successful at detecting focal PM targets, and focal PM responses incur lower costs on the OG task (leads to less increase in ongoing task response times). For instance, in the experiment described in Figure 1, it is easier to detect a PM target if it were *tortoise* as opposed to *tor*, since the full word *tortoise* is already being processed by the OG task. The focality effect interacts with the effect of emphasis, which we describe next (see Figure 2).

**Figure 2:**
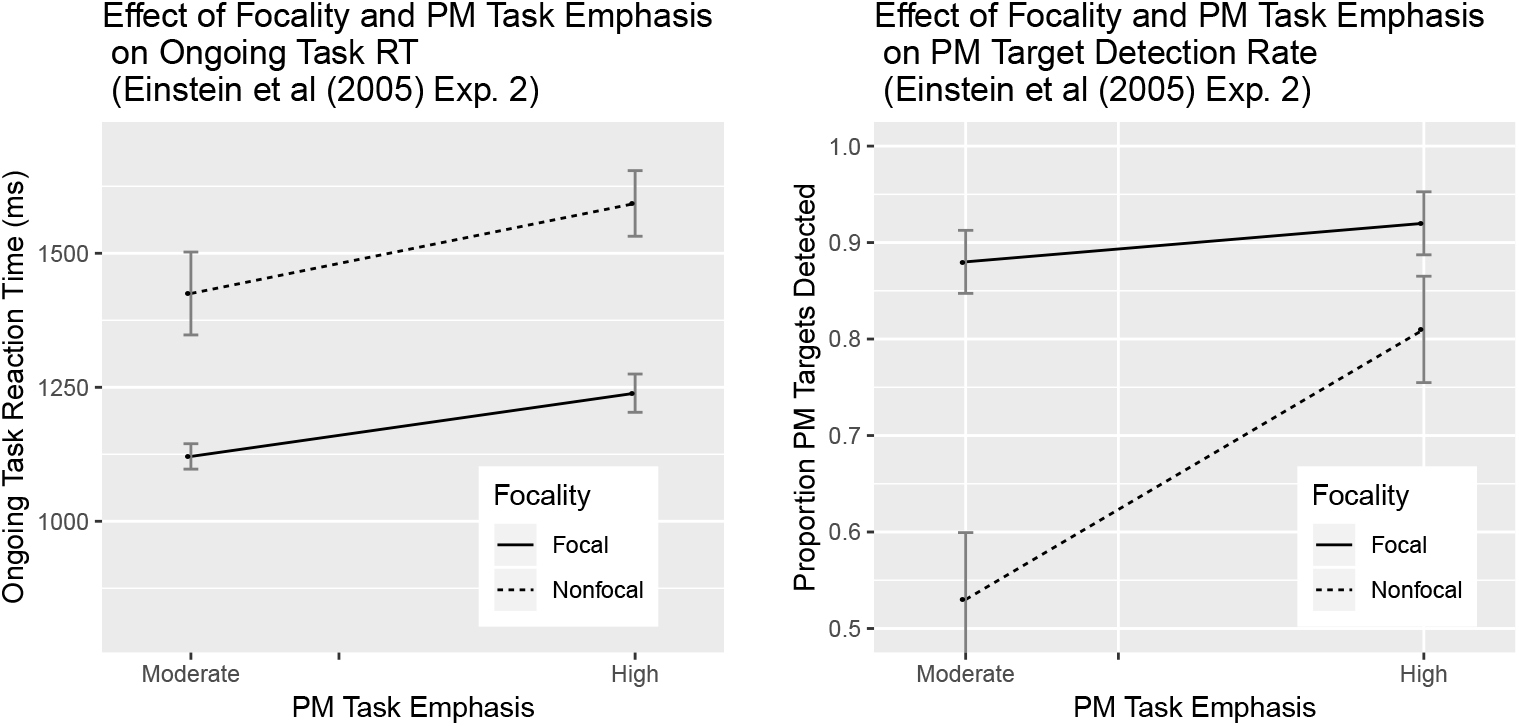
The interacting effects of *focality* and *emphasis* on ongoing task RTs and prospective memory hit-rate reported in Einstein et al. (2005). Performing an OG task takes longer when more emphasis is placed on the PM task. *Left:* A non-focal PM condition, where it is more difficult to detect the PM target, exerts a higher RT cost on the OG task. In contrast, in a focal PM condition, where PM target is more similar to OG task stimuli and easier to detect, exerts less of an OG task cost. OG task response times increase in both focal and non-focal conditions when high emphasis is placed on the PM task. *Right*: Participants exhibit high accuracy on focal prospective memory regardless of the emphasis condition. However, in non-focal conditions, emphasis has a large effect on the accuracy of PM responses.

#### Effects of relative task emphasis and interaction with focality

*Emphasis* refers to the relative priority that the participant is instructed to place on the PM vs. OG task. High emphasis indicates that the highest priority is to increase PM hit rate. For example, in the Einstein et al (2005) (Einstein and McDaniel, 2005) study, on half the trials the instruction emphasized the high importance of PM hit rate: *it is very important that you consider your main goal in this section to find absolutely every occurrence of the target item*. On the other half, PM hit rate was given moderate importance: *we have a secondary interest in your ability to remember to perform an action in the future*.

The main effect of emphasis is, sensibly, improved performance on the PM task as measured by PM hit rate. However, the instructional emphasis manipulation interacts with the focality effect (see Figure 2, lower panel): Emphasis has the largest effect on OG costs given non-focal targets (i.e., when PM targets are more difficult to detect because they involve detecting different features than those processed by the OG task). PM performance is at ceiling in the focal PM condition, when the two tasks share features.

In §4.2, we model emphasis by simply changing the ratio of reward associated with PM vs. OG task performance.

#### Effects of prospective memory load

The *load* or difficulty of the PM task has been empirically manipulated in a number of ways. In general, increasing PM load incurs greater costs on the OG task, measured by longer reaction times to ongoing task stimuli. For example, Einstein and colleagues showed that increasing the number of PM targets from 1 to 6 is associated with an increase in OG reaction times (Einstein et al., 2005) (Figure 4, left). Similarly, using a more difficult PM task (e.g., by increasing its working memory demand) also increases costs to OG reaction times (Meier and Zimmermann, 2015), and is shown to involve more anterior prefrontal cortex patterns of multivoxel fMRI activity (Momennejad and Haynes, 2013).

In §4.1 we model PM load in two ways: (a) by increasing the number of PM targets, and (b) by increasing the number of past, but no longer active, PM targets.

#### Individual differences and WM capacity

A number of studies have revealed systematic individual differences in prospective memory tasks. These differences are thought to be related to individual differences in working memory or executive function capacity. Brewer and colleagues Brewer and Marsh (2010) showed that individuals with lower measured WM capacity express a higher cost of PM on OG task response times in non-focal conditions. Furthermore, populations associated with lower WM or executive function capacities such as adolescents and older adults are more likely to benefit from strategies involving episodic memory, such as episodic future simulation, in PM performance (Altgassen et al., 2017, 2015). Individuals have also been shown to benefit from episodic encoding strategies such as imagery or implementation intentions (Brewer et al., 2011; Gollwitzer and Brandstätter, 1997).

In §4.3 we model individual difference effects by varying the process noise in the model, but retaining the assumptions of optimal control and evidence integration.

#### Strategy selection effects

As discussed in the Introduction, the multiprocess framework (Einstein et al., 2005; Einstein & McDaniel, 2005) suggests that there are two primary strategies participants use to perform the PM task. The first strategy involves active attentional monitoring for PM targets (presumably by maintaining this information in WM). The second strategy involves encoding the PM target in long-term memory, and relying on spontaneous associative recall of the PM task when the PM target appears. Several studies have sought to explicitly manipulate the choice between these strategies.

A series of studies controlled for strategy by encouraging participants to use episodic future simulation, imagery (Brewer and Marsh, 2010; Brewer et al., 2011), or an *implementation intention* strategy (Chen et al., 2015; Gollwitzer, 1990; McDaniel et al., 2008; McFarland and Glisky, 2012), in which they wrote down multiple times that upon seeing a PM target they will switch to the PM task. Varieties of episodic future simulation improved PM performance for non-focal PM. Crucially, the effects were eliminated if the subjects were not given a specific context for the associations. Furthermore, a recent study (Lewis-Peacock et al., 2016) directly manipulated the use of WM or episodic strategies in PM by either increasing proactive interference in episodic memory (hence increasing the benefits of a WM strategy) or increasing WM load by changing the ongoing task from 1-back to n-back (hence increasing the benefits of an EM strategy). Taken together, these studies help reveal the importance of the trade-off between WM and EM encoding strategies in PM performance.

The model’s strategy depends on the weight it gives to samples from WM and EM, and the optimal strategy is learned by the model given the task circumstances (e.g., high or load WM load) and the noise in the memory and perceptual samples.

## 3 A Rational Model of Episodic and Working Memory Use in Prospective Memory

We propose a simple rational model of the integrated use of noisy perception, working memory (WM) and longterm/episodic memory (LTM/EM) in service of prospective and ongoing tasks – as captured by the canonical dual-task paradigm described in Section 2.1. Our approach formalizes the multiprocess framework of PM proposed by Einstein and colleagues (Einstein et al., 2005) by specifying abstract computational properties of the LTM/EM, WM, and perceptual components, and asking how a model should use these components in the PM dual-task setting.

The model is *rational* (or normative) because the integration of evidence from the three components conforms to correct Bayesian inference, and because the policy (or strategy) parameters governing task performance are selected to maximize the joint reward on the OG and PM tasks. We also refer to it as *computationally rational* to emphasize that the model derives the best possible use of posited bounded computational resources Howes et al. (2009); it is thus an application of bounded optimality Russell and Subramanian (1995); Lewis et al. (2014). Once the (approximately) optimal policy parameters are computed and fixed (through a reinforcement learning method described below), the model provides detailed behavioral predictions, including response times and accuracies for both tasks. We explore these behavioral implications of the model in Section 4, compare them to human behavior, and discuss how the model explains the key phenomena summarized above.

### 3.1 Overview of the model’s processing in a single trial of the canonical dual-task paradigm

Before considering the mathematical details, it is useful to have a qualitative overview of how the three model components interact to perform one stimulus response in one trial of the event-based PM dual-task. We assume here that the ongoing (OG) task is a binary categorization task such as lexical decision, and the prospective (PM) task requires monitoring for a specific feature of the word (e.g., a syllabus). (Below we consider other variants such as monitoring for a set of words). We refer to the presented item as the *probe* item.

1. The *incentive reward* structure of the task is represented as a payoff matrix. This payoff matrix indicates gains and losses for making correct (or incorrect) responses to the OG task, for failing to make a response before a response deadline, and for correctly detecting (or missing) the PM target. These gains and losses can be used to establish some experimental conditions. For instance, PM emphasis can be determined by, or indicated in the payoff table as, the ratio of payoffs or gains for correct PM vs. correct OG performance. Thus, we can increase PM task *emphasis* by assigning a higher gain, or reward in the payoff table, for PM performance relative to OG performance.
2. Each stimulus item is represented as a feature pair: one feature is relevant for the OG task and one feature is relevant for the PM task. Feature values are drawn from a fixed set of discrete values. The OG task binary classification partitions the values into two subsets, one subset requiring a “Yes” response and one requiring a “No” response. The PM task requires matching the PM relevant feature to a previously presented PM target feature. The correlation between the two sets features can determine the *focality*. Thus, we can increase the degree of *focality* by increasing the correlation between the OG-relevant and PMrelevant features.
3. The model has a noisy encoding of the PM target(s) stored in episodic memory (EM). Each target encoding is represented as a discrete feature value which may be noisy, errorful with some small probability, and a discrete context code (indicating the PM trial) which may also be noisy and errorful with some small probability. In some model variants there may also be a corresponding encoding in working memory (WM). Note, here we indicate/increase *PM load* or *PM difficulty* conditions via two measures: (a) by increasing the number of PM targets that are monitored for simultaneously, or (b) increasing the number of past PM targets that are no longer relevant for the current trial.
4. The contents of EM (and possibly WM) and the task structure imputes a prior distribution, on possible stimulus probe items, in advance of any perception of the probe.
5. In this state, the model perceives the probe over discrete time steps. On each step, a noisy sample is drawn of the true pair of feature values representing the probe. Each sample step has a duration sampled from a gamma distribution.
6. On each noisy sample, the model updates a posterior distribution of possible target probes, and uses this distribution to compute a posterior over the three response hypotheses: the probability that the PM target-*yes* response is correct, the probability that the OG-*yes* response is correct, and the probability that the OG-*no* response is correct. This posterior *and* the time remaining until the deadline captures all the information the model needs to decide what to do next.
7. The noise in the sequential sampling process is an abstraction of multiple possible sources of process noise. This includes perceptual noise and noise in the integration of perceptual evidence with memory. We model *individual WM capacity* by changing this process noise level (Note that changing individual WM capacity is different from increasing WM load, e.g., via increasing the number of PM targets, described above.).
8. Given the posterior over correct responses and the time remaining until the deadline, the model chooses one of four possible actions: (1) obtain another noisy sample of probe features, (2) respond *yes* to the PM task (indicating that the model has detected the PM target), (3) respond *yes* to the OG task, or (4) respond *no* to the OG task. *The model makes this choice by choosing the action that maximizes expected payoff*. In an alternative task variant, the model must respond to OG task even when a PM target is present, and so the PM-*yes* action is replaced with two distinct actions, corresponding to PM-*yes*-OG-*yes* and PM-*yes*-OG-*no*.
9. The sum of the duration of probe sample steps before the choice is made constitutes a response time. The response type may be classified as a correct or incorrect OG response, a correct PM target detection, a PM target miss, a PM target false alarm, or a missed deadline.

In short, the model combines optimal evidence integration with optimal stopping and optimal response selection. Once memory and process noise parameters are fixed, we may obtain predictions of response times and accuracies on both PM and OG tasks. We do so under different task and architecture manipulations (e.g., PM emphasis, focality, PM difficulty, PM load, WM differences) by simulating the model many times. The primary technical challenges concern computing the prior distribution for probe items given the noisy memory encodings, and computing the policy for optimal stopping given the posterior and time remaining. We next present the formal model and describe how these two challenges are met.

### 3.2 Task environment and reward

#### Types of PM and OG tasks

There are two simultaneous tasks: the prospective memory (PM) task and the ongoing task (OG). *Probe* items are pairs of PM and OG features 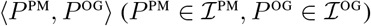 presented to the model for response before a deadline has passed There are *p* possible PM features 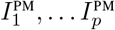 comprising the set 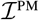, and *o* possible OG features 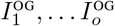 comprising the set 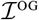. In our simulations, *p* = *o* = 10.

There are two kinds of PM tasks: detecting that a presented probe is the most recently presented target (where there may be a history of previous targets); or detecting that a presented probe is in a set of targets (there is no history of previous sets). Let 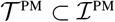 be the current set of PM targets.

There are two kinds of OG tasks: a binary discrimination (yes/no) of a presented probe (requiring no short-term memory); or detecting that a presented probe is the same as the probe presented 1 or 2 probes back (a working memory “N-back” task).

For the OG discrimination task, let 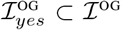 be the items that should elicit a *yes* response, and let 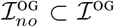 be the items that should elicit a *no*. 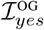 and 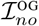 form a partition of 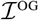.

There is a prior distribution of probe pairs that allows for a correlation *ρ*_Foc_ between OG and PM items/features. We discuss this below as a way to model *focality*.

#### Probe responses and task payoff

Consider first the case where the OG task is the discrimination task. Events of interest concerning the presented probes are:

1. PM_yes_: 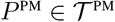 (probe is a/the PM target).
2. PM_no_: 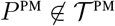 (probe is not a PM target).
3. OG_yes_: 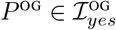.
4. OG_no_: 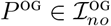.

So for given 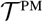, a probe 〈*P*^PM^, *P^OG^*〉 determines one of the three types of correct responses:

1. *Respond*-PM_yes_ should be made when PM_yes_, regardless of OG.
2. *Respond*-OG_yes_ should be made when PM_no_ and OG_yes_.
3. *Respond*-OG_no_ should be made when PM_no_ and OG_no_.

The fourth response is “no response”, which happens when the deadline is missed. There is a 4(*i*) x 3(*j*) task payoff matrix that indicates the payoff for making the *i*th response when response *j* is correct.

### 3.3 Noisy Encodings in Perception, EM, and WM

There are three cognitive components in the model: an episodic memory (LTM/EM) store, a strategically deployable WM, and noisy perception (Figure 3).

**Figure 3:**
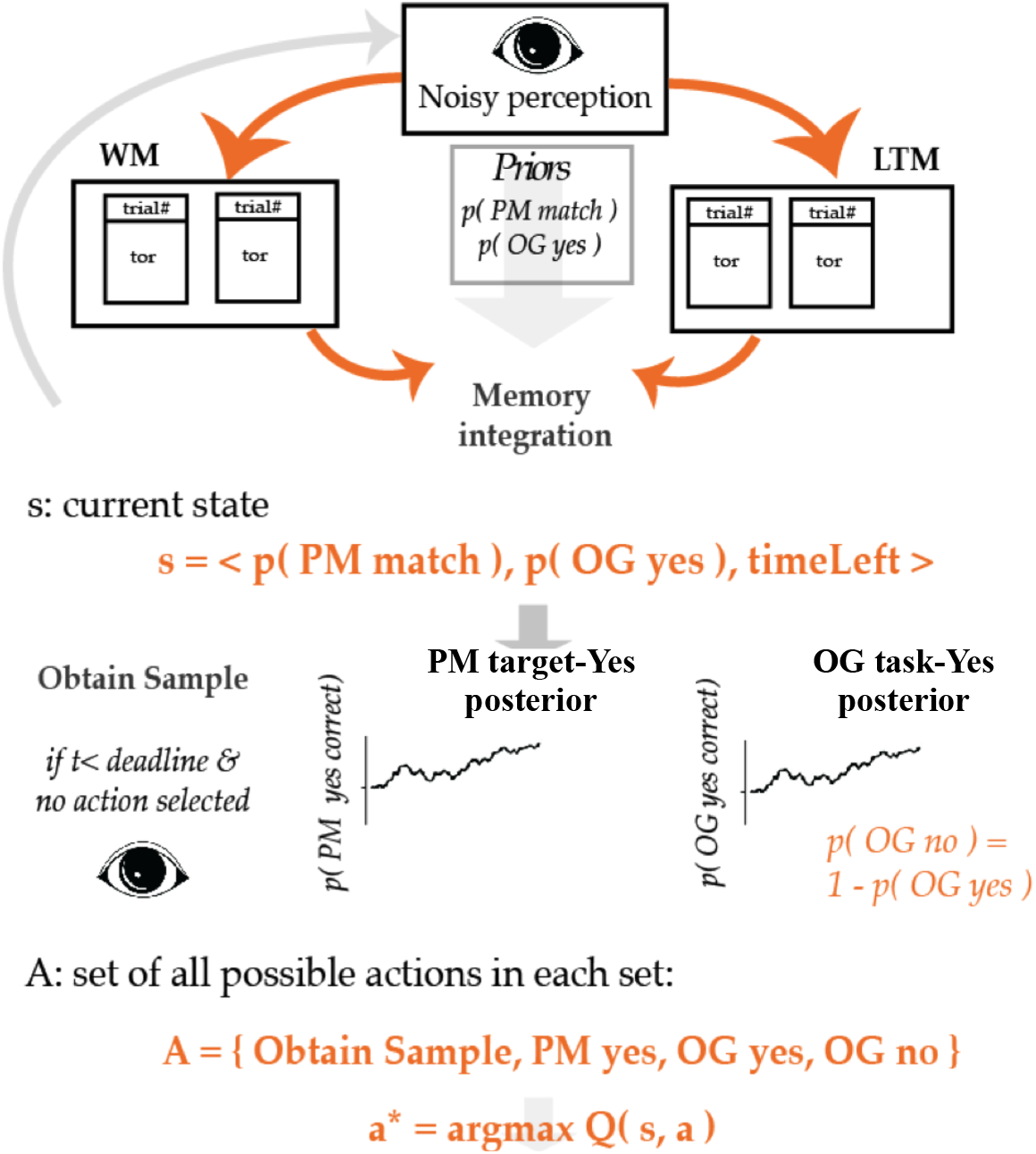
Model of rational WM-EM recruitment in a dual-task event-based prospective memory paradigm. The model components consist of a long-term episodic memory (EM/LTM) which stores a noisy encoding of current and past PM targets, and a working memory (WM) in which the ongoing stimulus and PM target are encoded with some noise. Parallel accumulators with no bounds draw noisy perceptual samples of the OG and PM features of the probe stimulus, incrementally updating posteriors that track the status of the probe for the PM task and the status of the probe for the OG task. For every state, the set of all possible actions includes obtaining another sample (waiting longer), giving a PM *yes* response, an OG *yes* response (e.g., category match or 1-back match), or an OG *no* response. The optimal policy is computed using Q-learning (Watkins and Dayan, 1992), which selects the action with the highest expected value. At each time point the optimal policy determines whether to draw another sample and risk going past deadline, or to make a response. Bayesian integration weighs information in WM and LTM/EM as a function of the uncertainty of the memory encodings.

#### The LTM/EM encodings

EM is a set of elements that are noisy encodings of current or past PM targets in long term or episodic memory. Each encoding consists of a PM feature value and a context feature value. The PM feature value may be noisy and erroneous: with some small probability, the feature is drawn from other distractor feature values rather than the correct feature value. Context codes represent the trial number of past PM targets. Context codes may also be erroneous: with some small probability, a context code is drawn from context codes for other trials; we assume a similarity confusion matrix for trial codes such that nearby trials are more confusable.

#### WM encodings

WM is a set of elements that are noisy encodings of OG task stimuli and/or the current PM target. When the OG task is N-back, as in (Lewis-Peacock et al., 2016), WM contains the past N-back (1 or 2) stimuli and the current PM target (depending on task demands and strategic choices, as described below). While we do not model experiments with N-back in the present manuscript, the model is theoretically reach enough to be applied to these studies as well.

Context codes in WM can, for instance, represent the N-back position of an item (e.g., 1 back or 2 back). In the N-back=1 condition, the most recent N-back probe is kept in WM. In the N-back=2 condition, the two most recent probes are kept in WM. If a WM encoding strategy is used for PM, the PM target encoding is also present in WM. The item and context codes are not confusable across PM targets and N-back stimuli, but increasing the number of items held in WM is assumed to increase the item and context noise.

#### Perceptual samples and processing noise

The true probe identity is a pair of feature values 〈*P*^PM^, *P^OG^*〉. When the model perceives the probe, with some probability *ϵ*_Proc_ the PM feature of the perceptual sample is picked uniformly from non-true values of PM feature, otherwise it is the true feature. The same is done independently for the OG feature of the stimulus. We refer to this noise as *processing noise* (rather than perceptual noise) to emphasisze that it an abstraction over multiple sources of noise, including perceptual or attentional noise and noise in the evidence integration process.

### 3.4 Bayesian integration of noisy evidence in memory with noisy perceptual samples

Before any perceptual samples of the stimuli have been obtained, the noisy WM and EM memory encodings along with information about the base rate of PM target occurences are used to compute expectations: a prior distribution over possible PM targets and OG stimuli. This prior distribution is updated as noisy perceptual samples are obtained.

From this distribution a posterior over three response hypotheses is computed: the posterior probability that the PM target-*yes* response is correct, the posterior probability that the OG-*yes* response is correct, and the posterior probability that the OG-*no* response is correct. More formally:

#### Desired response posteriors given samples

When the model arrives at a probe 〈*P*^PM^, *P^OG^*〉 with some memory *m*, it sequentially draws perceptual samples of the presented probe, integrates the evidence, and makes a response after some number of samples are drawn (or fails to respond if a deadline has passed) as follows.

Here a sequence of *k* perceptual samples of the true PM feature *P*^PM^ are denoted as 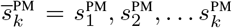 and the sequence of perceptual samples of the true OG feature *P*^OG^ as 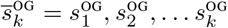.

After *k* samples, the model computes the posteriors for each of the three responses being the correct response:

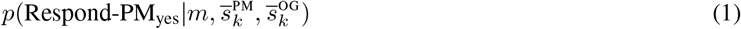

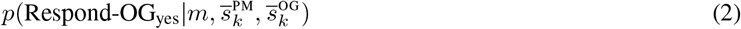

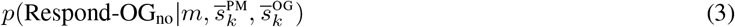

Given these posteriors, the expected payoff of the three responses may be computed from the payoff matrix, and compared to the expected value of obtaining another sample (how this value is estimated is described below).

The posteriors over the three responses may be computed as a function of the four probe matching cases of (PM_yes_, PM_no_, OG_yes_, OG_no_):

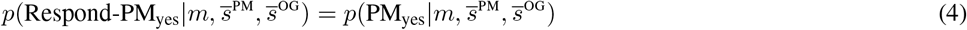

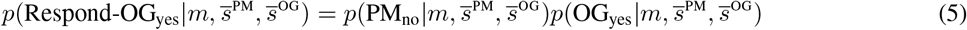

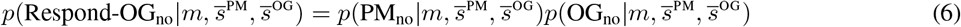

These three equations capture the task instructions concerning responses. Next we describe how the probe-match posterior probabilities are sequentially computed as each sample arrives.

#### Posterior update given perceptual samples

The perceptual samples are not conditionally independent given the abstract event types (PM_yes_, PM_no_, OG_yes_, OG_no_) because they depend on the specific presented probe pair. The samples *are* conditionally independent given a specific probe pair, because the noise associated with perceptual samples is uncorrelated across time. We exploit this conditional independence to allow for an incremental update of the posterior given each perceptual sample.

The specific probe pair is unknown to the model (hence the need for the perception). We handle the dependence of the posterior quantities in Equations (1)–(3) on the previous samples (and on the memory *m*) by updating a distribution over probe pairs.

Consider the update for PM_yes_ given a perceptual sample *s*^PM^:

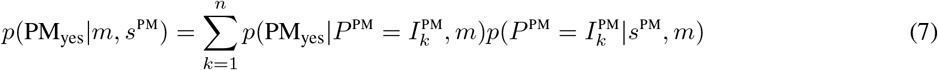

We must now compute the two terms in the product on the righthand side. 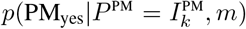 is the probability of a PM target match, given that the presented probe is 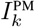 and the model has memory *m*.

Intuitively, each different state of the memory determines a different relationship between presented probe and desired response. This quantity is computed from a joint probability table estimated by large scale Monte Carlo simulation.

The other term 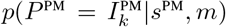 is the posterior over 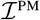 given the perceptual sample, and is computed with Bayes rule:

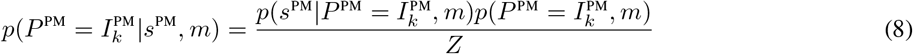

where 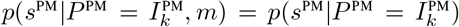 is the perceptual noise model and *Z* is the normalization term. The term 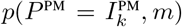 captures the dependence of the probe on the agent’s noisy memory, which may include multiple (noisy) PM targets and WM encodings. It is not possible to derive this quantity analytically, but precise estimates may be computed from empirical frequencies obtained via large scale Monte Carlo simulation.

### 3.5 Computing the optimal policy via reinforcement learning

At each time step, the model chooses among four possible actions: *Respond*-PM_yes_, *Respond*-OG_yes_, *Respond*-OG_no_, and *obtainSample*.

After obtaining *k* samples, the model calculates the expected value of *Respond*-PM_yes_, *Respond*-OG_yes_ and *Respond*-OG_no_(via the posteriors (1)–(3) and the payoff matrix). The model must compare these expected values to the value of the obtain another sample action (*obtainSample*). This value depends on the time remaining *t_rem_* before the deadline and the current belief (uncertainty) concerning which experiment state (PM_yes_, etc) holds. The belief is fully captured by the posteriors *p*(PM_yes_|*m*, *s*^PM^) and *p*(OG_yes_|*m*, *s*^OG^). The state *s* for conditioning control is therefore the triple

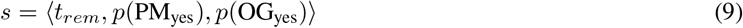

Thus, by computing the optimal value function *Q*\* (*s*, *obtainSample*) for all model states, we can simulate the rational model behavior for any set of experimental trials. We estimate this value function by using tabular Q-learning with a discrete binned approximation to the value function: each of the three continuous state variables is mapped to one of *b* bins, where *b* is a hyper-parameter of the learning. Computational experiments show best performance around *b* = 50 (dependent of course on other hyper-parameters). The Q-learner need only learn the value of the action *obtainSample*, because the values of the other three actions (the three task responses) can be computed directly from the posteriors at each time step. No temporal discounting is used to define expected value.

## 4 The Model’s Account of the Key Phenomena

We used the model described in Section 3 to simulate a number of behavioral phenomena in the human PM literature outlined in Section 3. Most of these phenomena involve two behavioral measures of interest: reaction times to the OG task and the accuracy (detection rate) of responses to the PM target.

All simulations use a single consistent setting of memory noise parameters. We set *task* parameters, including payoff, to plausible values that approximate a canonical PM paradigm (Einstein et al., 2005). We set the *agent* process and memory noise parameters to values that yield human-level accuracies on the PM and OG tasks. Note that these parameters are not adjusted per condition to match the empirical phenomena.

Table 1 summarizes these parameters. The payoffs for the PM target detection are relative to a fixed value of 1 (−1) for a correct/incorrect OG task response. The range of PM payoffs are higher (10–30) because PM target probes appear much less often that OG probes, and thus higher PM payoffs are needed to create overall payoff balance between the two tasks.

**Table 1:** Task and agent parameters used in the simulations.

|  | Parameter | Description | Value(s) |
| --- | --- | --- | --- |
| TASK | $R_{OG}$ | Reward/payoff (penalty) for successful OG response | 1 (-1) |
| | $R_{PM}$ | Reward/payoff (penalty) of successful PM target detection | 10 (-10); 15–30 for higher PM emphasis |
| | $R_{FA}$ | Penalty for PM target false alarm | -10 |
| | $p_{target}$ | Probability of PM target occurrence | 0.05 |
| | $\rho_{Foc}$ | Focality: correlation of PM and OG features | 0–0.2 (non-focal); 0.8–1.0 (focal) |
| | $d$ | Deadline | 100 |
| AGENT | $\epsilon_{Proc}$ | Process/sample noise (error rate) | 0.4 (0.2 for high-WM capacity) |
| | $\tau_{Mean}$ | Mean sample duration | 25 |
| | $\tau_{CV}$ | Sample duration coef. of var. | 0.3 |
| | $\epsilon_{EM}$ | EM/LTM noise (error rate) | 0.002 |

### 4.1 Effect of PM load on prospective and ongoing task performance

As noted in §2.2, one of the most stable and widely reported effects related to PM is the cost of a PM task on the reaction times to the OG task (Einstein and McDaniel, 2005; Meier and Zimmermann, 2015; Pink and Dodson, 2013). That is, performance of an OG task is slower in the presence of a prospective memory intention, and this cost increases with PM load—for example, the number of PM targets (Einstein and McDaniel, 2005) or the demands that these place on WM (Momennejad and Haynes, 2013; Meier and Zimmermann, 2015; Lewis-Peacock et al., 2016).

Furthermore, the cost of PM is exacerbated when higher emphasis is placed on the PM task, which we have here operationalized as higher priority or reward for correct PM responses, e.g., priority or importance manipulation (Einstein and McDaniel, 2005). Finally, manipulations that favor the use of WM to perform the PM task increase PM costs as well (Einstein and McDaniel, 2005).

Figure 4, right, shows the model’s simulated reaction times to the ongoing task as a function of the presence of the PM task, PM load as manipulated by changing the number of PM targets, and emphasis on the PM task. The model shows a clear effect of PM task presence and load: responses to the ongoing task are slowed, and this slowing increases when higher emphasis is placed on the PM task (which is here operationalized as increased PM task payoff relative to OG task payoff).

**Figure 4:**
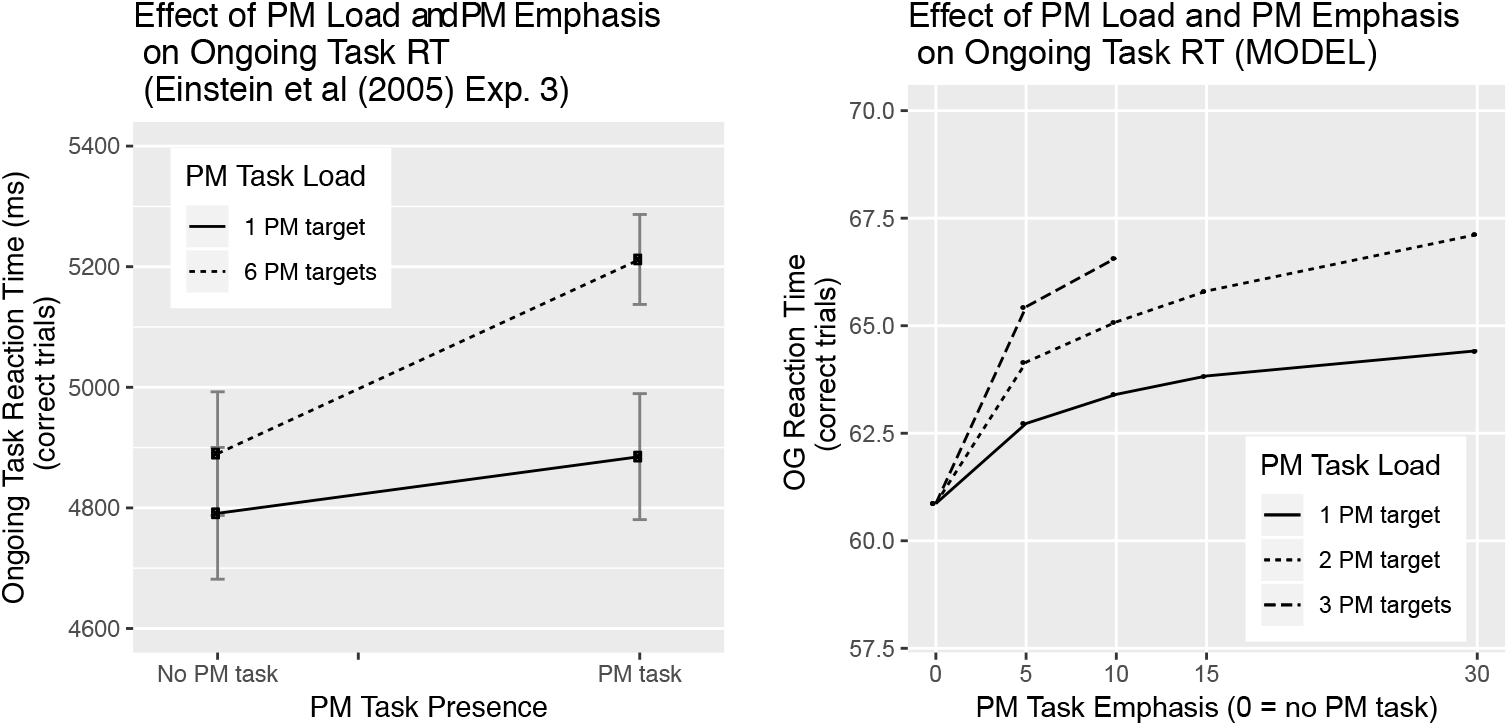
Effect of PM load on ongoing task RTs. We manipulated PM load by changing the number of targets and observed its effect on OG reaction times (as Einstein and colleagues did) and simulated its interaction with PM emphasis. Human behavior (left) and model simulation (right) are shown here. The presence of a PM task slows down performance on the OG task. This slowing effect is exacerbated by an increase in the number of PM targets. The increased number of samples before giving a response corresponds to an increase in OG reaction times. The behavioral experiment (left) compares the RT costs of 1 vs. 6 PM targets (in the presence of either of these targets the participant must perform the PM task) for the OG task. The model (right) qualitatively simulates the effect by comparing 1, 2 and 3 PM targets. Model predictions (right, Y axis unit: #samples taken before response) qualitatively reproduce the human behavioral findings from Einstein et al. 2005 (left, Y axis unit: seconds).

The model derives this slowing on the OG task as the *computationally rational response* to the changes in the task parameters. What has changed is the optimal stopping criterion (derived via Q-learning). In the presence of the PM task, the agent faces a different response discrimination demand: not only is the agent discriminating among the OG features but it is, in parallel, discriminating among the PM features, and additional samples are required (in expectation) to reduce uncertainty about the status of the probe.

This dependence of the OG response on PM certainty is captured in Equations (2) and (3). Increasing the PM load by increasing the number of targets further slows the OG task. In followup simulations we confirmed that this slowing is not merely due to the fact that each individual target has a lower probability of occurrence.

### 4.2 Effect of focality and emphasis

In non-focal PM, the PM targets demand attention to other features than those probed by the OG task. For instance, the OG task might require a category match or 1-back word match judgment, whereas the PM task may depend on specific syllables or the font color. A robust finding in the literature is that human participants display higher reaction times to the OG task in the non-focal PM condition, see Figure 6, left (Einstein and McDaniel, 2005).

We model *focality* by manipulating the correlation between the OG and PM features. The intuition is that focal PM features are highly correlated with OG features, while non-focal PM features are not correlated with OG features. Put differently, in focal conditions, OG task features also provide useful information about PM target status. The correlation parameter *ρ*_Foc_ thus provides a way to continuously vary focality, from extremes of 0 (non-focal) to 1 (focal with complete overlap of features). Many tasks in the literature are sufficiently complex that is not possible to quantiatively estimate what this parameter should be in order to simulate specific experiments (e.g., the features of the word ’tortoise’ and syllable ’tor’ may be more correlated than the features of a word to the font color). Thus, here we explore a range of low and high values to assess the qualitative predictions.

*Emphasis* manipulations have been realized experimentally through changes in instructions; as described above. We model emphasis changes by increasing the payoff and penalties (relative to the OG task) for succcesful detection and misses for the prospective memory task. Because the experimental manipulations are instructional, it is not possible to precisely estimate what the PM payoff should be. We explore here a range of PM payoffs that change the total proportion of reward obtained from moderate level (about 1/4 of the total reward due to the PM task) to relatively high (about 2/3 of the total reward obtained due to the PM task).

As shown in Figure 5, the model qualitatively reproduces the effects of both focality and emphasis and their interaction. Thus, the computuationally rational response—in the precise sense of maximizing expected task payoff given the computational constraints—to both non-focality and higher PM empahsis is to slow down.

**Figure 5:**
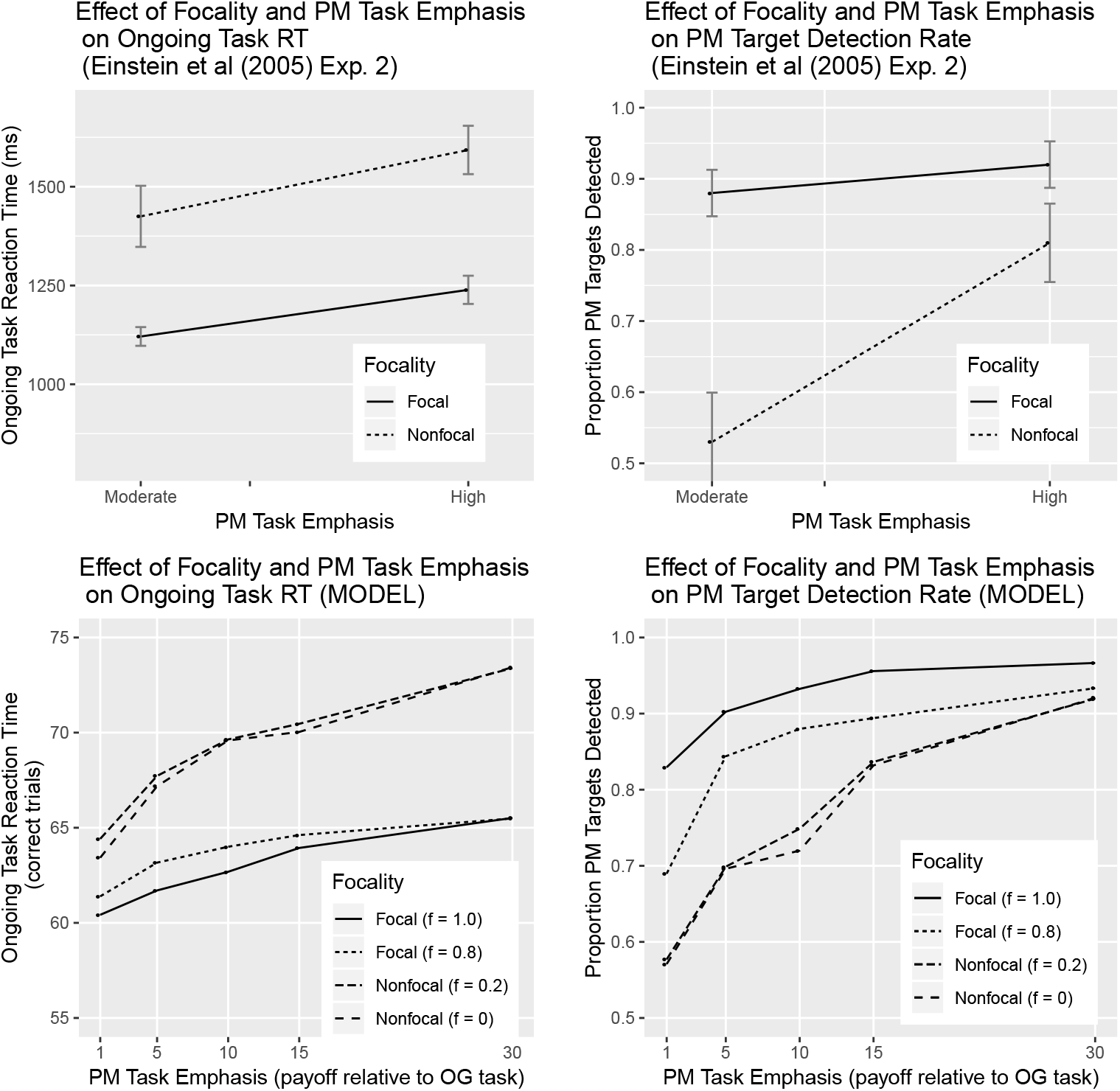
The model produces focality and PM emphasis effects. Human results (see Figure 2) and corresponding model simulation results (bottom). Both humans and model simulations show higher costs in the non-focal compared to the focal condition. This effect is modulated by emphasis on the prospective memory task. While human experiments only included high and low emphasis instruction effects (two conditions), the model simulates the effect of both emphasis and focality along continuous dimensions.

Under conditions of high focality, the information obtained from both OG and PM features is yoked. Therefore, when information has accumulated quickly for the OG task, it has also accumulated quickly for the PM task; thus rapid OG responses need not be delayed to wait for the PM target discrimination. However, as PM emphasis (relative PM payoff) increases, the gains in the expected value associated with greater PM discrimination certainty outweigh the risks of missing the deadline. Thus, given high PM emphasis, OG responses are slowed down until PM discrimination reaches higher certainty.

There is a complex tradeoff and interplay among all these factors—deadline risk, OG payoff, PM payoff, correlation of PM and OG features—but optimally navigating this tradeoff and complexity is precisely what the rational evidence integration and optimal control policy does.

### 4.3 Effect of individual differences in working memory capacity

Brewer et al (2010) Brewer and Marsh (2010) explored the role of individual cognitive differences by administering two standard span measures of working memory capacity to participants, and separately examining the performance of those who scored high and low on the composite WM capacity measure. They showed that participants with lower measured WM capacity displayed a large effect of focality on both PM target detection accuracies and PM target detection reaction times, while high working memory capacity participants showed little effect of focality (Figure 6, top row, left and middle). Both groups showed increased reaction times to the OG task in non-focal conditions and there were no overall difference in reaction times between the two groups.

**Figure 6:**
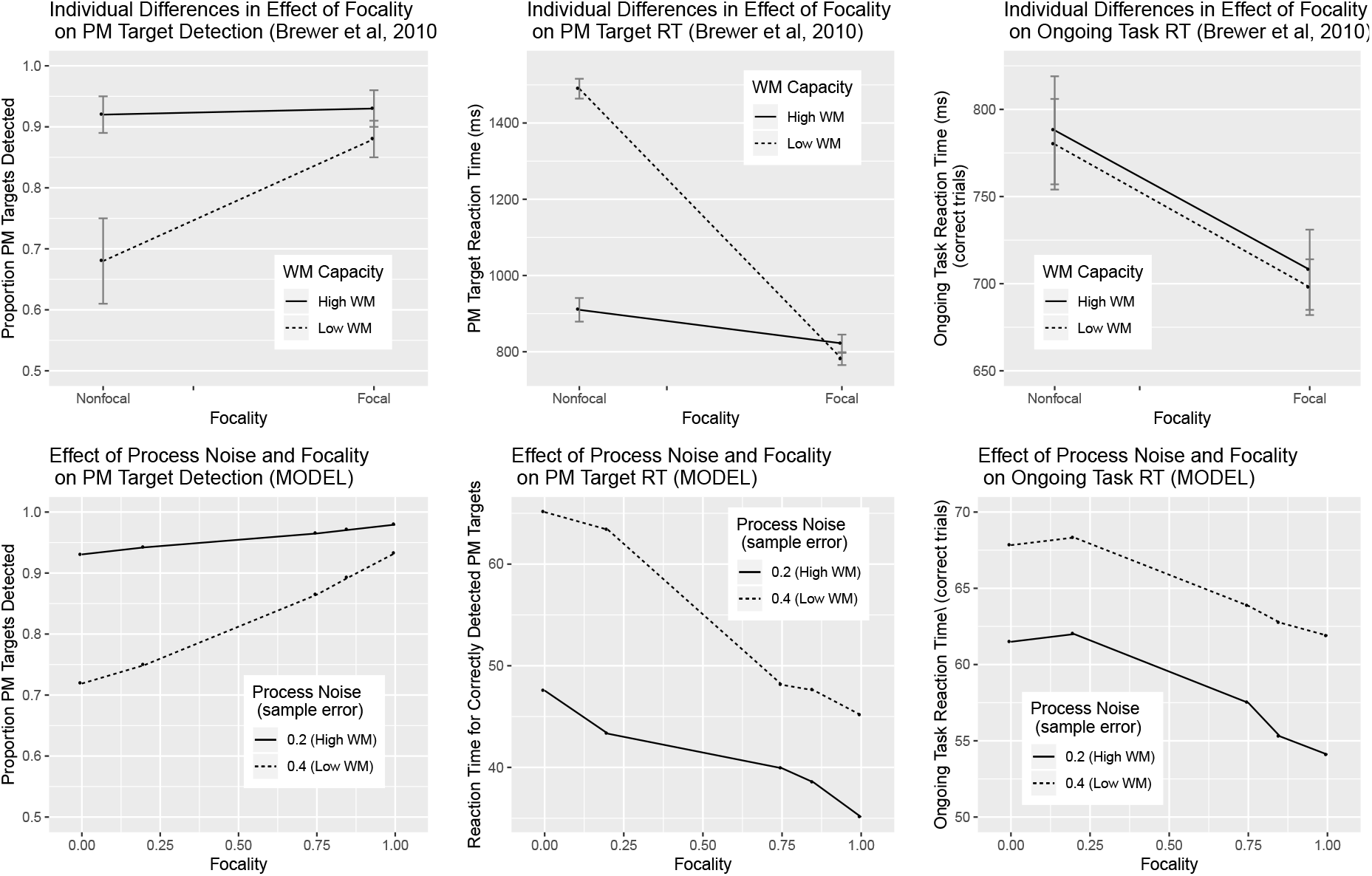
Effect of individual differences on OG and PM performance. Brewer et al (2010) found robust differences in performance on PM detection and rseponse time between participants with low vs. high measured working memory capacity (top row). By changing the process (sampling) noise parameter, the model accounts for these differences in both PM detection rate and RT, and their interaction with focality (bottom row). The model correctly predicts no interaction between focality and WM capacity in OG reaction times (right), but incorrectly predicts an ovderall difference in OG RT not observed in the Brewer et al (2010) task.

We modeled WM capacity differences by manipulating the process noise (sampling error); in this model lower process noise corresponds to higher working memory capacity.

This manipulation accounts for both the PM target detection accuracy and PM target response time interactions between focality and working memory capacity (Figure 6, bottom row, left and middle). The overall effect of focality on OG task reaction time, and the lack of an interaction with working memory capacity, is also accounted for.

However, the model incorrectly predicts a main effect of working memory capacity on RT (predicting faster responses for higher WM), an effect not observed by Brewer et al. It is possible that the two particpiant groups also differ in subjective payoffs for speed and accuracy, or relative emphasis on the PM task, all of which could diminish the predicted difference on OG RTs between the two groups, but our intent here was to understand what could be accounted for by varying only a single parameter.

### 4.4 Clarification of key distinctive properties of the rational model

We now summarize some of the key properties of the model that distingish it from other approaches.

1. *Value maximization and evidence accumuation without response thresholds*. There are no thresholds or bounds for the evidence accumulation. Rather, for every given state, defined in terms of 〈*t_rem_*, *p*(PM_yes_), *p*(OG_yes_)〉, a response is selected once the expected value of giving one of the responses is higher than the expected value obtaining another sample. In other words, the *stopping policy space* here is the full policy space conditioned on the two posteriors and time remaining. Any policy space with thresholds is a strict subset of this space.
2. *Optimal weighting of information in EM, WM and perception via Bayesian integration*. The Bayesian integration naturally weights information in WM and LTM/EM as a function of the uncertainty of the memory encodings. When LTM/EM noise or interference is high, e.g., due to the presence of lures, relatively lower weight is given to LTM/EM. When information in WM is noisier (e.g., due to high load), relatively lower weight is given to WM. There are no explicit weight parameters; rather weighting is a natural consequence of the likelihood function that captures the dependence of the prior on noisy memory.
3. *Use of noisy information in memory without a separate retrieval process*. The model contains no explicit retrieval processes; rather, the model conditions its responses on all the information in the noisy memory store. This global-parallel property is consistent with most mathematical models of memory retrieval that assume parallel contact with all memory elements (e.g., (e.g., Shiffrin, 2003)). But the model provides accounts of reaction times and error rates without assuming a retrieval process that yields an item or subset of items from memory that must then be further processed.
4. *Sensitivity to differential task emphasis and focality manipulations without explicit resource or attention allocation mechanisms*. The model provides accounts of task emphasis and focality effects through purely task-environment manipulations (changing payoff structure and probe-feature correlations) and without recourse to any attention or resource allocation mechanism. Because these effects are robust consequences of the task environment manipulations, the rational model provides a useful baseline against which predictions of resource allocation mechanism theories may be compared. Under the present account, focality and emphasis effects are not signatures of adaptive resource allocation, but signatures of a rational stopping criterion given noisy memory stores.

## 5 Summary and future directions

Here we combine Bayesian inference and reinforcement learning to offer a normative solution to the prospective memory problem. The prospective memory problem is that of the simultaneous and timely execution of immediate (ongoing) and delayed (prospective) tasks. We have proposed a tripartite model of prospective memory function with noisy perception and processing, episodic memory, and working memory components (Section 3). The model is rational and normative because the integration of evidence from the three components conforms to correct Bayesian inference, and because the policy parameters governing task performance are selected to maximize payoff on both the ongoing and prospective tasks (using RL).

The model is *computationally rational* because it derives the best possible use of posited bounded computational resources Howes et al. (2009); Lewis et al. (2014). Once the (approximately) optimal policy parameters are computed (here through reinforcement learning), the model provides detailed behavioral predictions, including response times and accuracies for both ongoing and prospective tasks. The model thus provides a formally rigorous account for understanding interactions between long term and working memory in the service of prospective memory.

A promising future direction is the application of the model to existing neuroimaging data on prospective memory. Over the past decade, a number of fMRI studies have shown that univariate as well as multivariate patterns of activations in the prefrontal cortex, the parietal cortex, and the hippocampus mediate event based and time-based prospective memory (Gilbert, 2011; Momennejad and Haynes, 2012, 2013), as well as the interaction of LTM/EM and WM processes in successful PM (Lewis-Peacock et al., 2016). Future studies could study the distinct and joint computational processes governing the prefrontal components (often involved in WM and controlled processing) and the hippocampal components (involved in associative memory, LTM/EM) of these findings. Of particular interest would be to compare the function of these regions in healthy and abnormal PM behavior in order to identify predictors of the optimality of action selection in the model.

A critical contribution of the model, in all of these contexts, is a normative account of the strategic gradient in the deployment of LTM/EM and WM in the performance of memory-dependent tasks. Such a normative model addresses the following question: how should LTM/EM and WM be used to realize planned action? As such, the model provides a valuable foundation for understanding interactions between LTM/EM, WM, and perception in other multitasking settings. This may be useful in understanding the meta parameters underlying the performance of healthy individuals, and accordingly designing models that solve real-world multi-tasking problems.

Such meta parameters may also allow us to understand the bounded rationality of parameters that may compromise function due to brain injury or psychiatric conditions. Many psychiatric disorders are marked by impairments in the adaptive integration of perception with multiple memory systems. By providing insights into underlying mechanisms of normal and suboptimal adaptive behavior, our model could be used to design compensating interventions in participants whose behavior suggest sub-optimal integration of WM and EM sources of memory. Such computational interventions could help improve real-world performance in both healthy individuals and those in sub-optimal conditions.

